# Multi-axis spatiotemporal niche partitioning between coexisting top predators in ponds

**DOI:** 10.64898/2026.01.09.698714

**Authors:** Jason R. Rohr

**Affiliations:** Department of Biological Sciences, University of Notre Dame, Notre Dame, IN 46556, USA

**Keywords:** activity patterns, amphibians, behavioral thermoregulation, coexistence, diel activity, freshwater ecology, habitat partitioning, niche overlap, predator–predator interactions, spatiotemporal niche partitioning, temperature dependence

## Abstract

Understanding how ecologically similar species coexist remains a central challenge in ecology, particularly in small, spatially constrained systems where opportunities for segregation may be limited. Classical niche theory predicts that coexistence is facilitated when species partition resources across multiple niche axes, yet few empirical studies quantify how spatial, temporal, and environmental dimensions jointly structure realized niches in natural systems.
We examined spatiotemporal niche partitioning between two coexisting top predators—the eastern red-spotted newt (*Notophthalmus viridescens*) and bluegill sunfish (*Lepomis macrochirus*)—in a pond lacking piscivorous fish. Using year-round trapping data collected across depth strata, diel periods, and seasons, we combined hierarchical count models, temperature-informed partial effect analyses, and null-model tests of niche overlap.
Newts and sunfish exhibited strongly contrasting patterns of habitat use across multiple axes. Newt capture rates were highest during cooler periods, in deeper habitats, and during morning sampling, whereas sunfish capture rates peaked during warmer periods, in shallow habitats, and during afternoon sampling. Model-based analyses revealed opposing responses to temperature, with predicted newt captures declining and sunfish captures increasing as temperature rose, even after accounting for seasonal effects. As a result, niche overlap across combined season-by-depth-by-time states was consistently lower than expected under randomized null models.
Across all analyses, newts and sunfish exhibited strong and consistent spatiotemporal niche partitioning, with opposing seasonal trajectories, contrasting depth and diel activity patterns, divergent thermal responses, and niche overlap significantly lower than expected under null models. These results demonstrate that fine-scale spatiotemporal structure across interacting niche axes can generate pronounced segregation among coexisting top predators, even in small and physically constrained ecosystems. Rather than reflecting partitioning along a single dominant axis, niche differentiation in this system appears to emerge from the coordinated interaction of season, habitat, diel activity, and temperature, highlighting how multi-axis dynamics shape realized niches in natural communities.

**Significance Statement:** Coexisting predators often exploit the same prey and habitats, raising the question of how overlap is reduced in spatially constrained ecosystems such as ponds. By integrating seasonal, diel, habitat, and thermal dimensions, this study demonstrates that two ecologically similar top predators—newts and sunfish—exhibit strong spatiotemporal niche partitioning that substantially lowers overlap relative to random expectations. Our results show that fine-scale temporal and habitat structure can play a major role in organizing predator assemblages, even in small freshwater systems, and highlight the importance of multi-axis niche frameworks for understanding species interactions and persistence in natural communities.

## INTRODUCTION

Understanding how ecologically similar species coexist remains a central goal of ecology. Classical niche theory predicts that coexistence is promoted when species differ in their use of limiting resources across one or more niche axes, thereby reducing direct competition (Hutchinson 1957; Schoener 1974). Empirical studies have documented niche partitioning along spatial, temporal, and trophic dimensions across a wide range of taxa (Schoener 1974; Chesson 2000; Barabás, D’Andrea & Stump 2018; Ellner *et al*. 2019). However, the relative importance of these axes, and the extent to which they interact, often remains unresolved, particularly in natural systems where environmental conditions vary predictably over diel and seasonal timescales (Bloomfield *et al*. 2022).

Recent and historical work has emphasized temporal niche partitioning as a key mechanism structuring ecological communities, including diel activity segregation, seasonal phenology, and timing of habitat use (Schoener 1974; Kronfeld-Schor & Dayan 2003; Carscadden *et al*. 2020; Cox & Gaston 2024). Many empirical studies have demonstrated temporal segregation among species, yet these studies frequently focus on a single temporal dimension (e.g. diel or seasonal activity), rely on indirect proxies of activity (such as detection times, stomach contents, or telemetry records), or compare populations across sites rather than among coexisting species within the same system (Kronfeld-Schor & Dayan 2003; Rowcliffe *et al*. 2014; Bloomfield *et al*. 2022; Cox & Gaston 2024). Consequently, although temporal niche partitioning is well documented, fewer studies quantify how diel, seasonal, and habitat dimensions jointly structure realized niches in situ, particularly while accounting for environmental drivers, such as temperature, that impose strong physiological and behavioral constraints.

Small freshwater ponds provide tractable systems for examining spatiotemporal niche structure among consumers because physical gradients are steep, habitats are shared, and species interactions can be strong. In ponds free of piscivorous fish, the coexistence of eastern red-spotted newts, *Notophthalmus viridescens* Rafinesque, and bluegill sunfish, *Lepomis macrochirus* Rafinesque, represents a case of co-occurring, ecologically similar top predators. Both species function as keystone predators (sensu Paine 1969) because they disproportionately eat invertebrate consumers and competitively dominant prey, thereby exerting strong top-down effects on freshwater communities (Morin 1981; Crowder & Cooper 1982; Morin 1983; Gilinsky 1984; Morin 1984b; Morin 1984a; Butler 1989). Although dietary overlap between newts and sunfish is substantial (Butler 1989; but see Kurzava & Morin 1998), relatively little is known about the extent to which these species partition other major niche axes, particularly habitat use and temporal activity, despite the potential for strong constraints imposed by predation risk, diel cycles, and seasonal temperature variation.

In addition to representing a system of coexisting, ecologically similar predators, the newt–sunfish interaction offers a relatively rare example of an amphibian species coexisting with fish (Kats, Petranka & Sih 1988; Wellborn, Skelly & Werner 1996). Adult *N. viridescens* often perform better in the absence of sunfish (Bristow 1991; Hecnar & M’Closkey 1996), suggesting that fish may exert competitive or behavioral constraints on newts. Experimental work has shown that sunfish can outcompete adult newts under warm conditions (Bristow 1991), highlighting the potential for environmental context, particularly temperature, to mediate interactions between these species. Nevertheless, newts and sunfish commonly coexist in natural pond systems, indicating that processes operating across spatial or temporal dimensions may facilitate their persistence.

We hypothesized that coexisting newts and sunfish exhibit spatiotemporal niche partitioning across diel, seasonal, and habitat axes, consistent with differences in thermal ecology and activity patterns. Specifically, we predicted that (i) relative capture rates of newts would be concentrated in cooler conditions, deeper habitats, and morning periods, whereas sunfish would be concentrated in warmer conditions, shallower habitats, and afternoon periods (Schoener 1974; Werner & Hall 1988); (ii) seasonal shifts in temperature would be associated with opposing changes in capture rates of the two species, reflecting differential responses to thermal conditions (Huey & Kingsolver 1989; Angilletta 2009); and (iii) overlap in resource use across combined season-by-depth-by-time states would be lower than expected under null models of random habitat and time use (Pianka 1973; Gotelli & Graves 1996).

To evaluate these predictions, we used year-round trapping data and combined hierarchical count models, temperature-informed partial effect analyses, and null-model tests of niche overlap. Rather than testing whether niche partitioning causes coexistence, our goal was to quantify the structure and strength of spatiotemporal niche differentiation among coexisting top predators and assess whether these patterns are consistent with stabilizing niche differences predicted by coexistence theory.

## MATERIALS AND METHODS

### Study system and field sampling

The study was conducted in the piscivorous fish–free side pool of Harpur Pond (Binghamton University, Broome County, New York, USA). This pond is typical of many permanent ponds in the northeastern United States and has a maximum depth of approximately 1.33 m at 3–5 m from shore. The pond supports heavy beaver activity and is surrounded by mixed deciduous forest. Eastern red-spotted newts (*Notophthalmus viridescens*) and bluegill sunfish (*Lepomis macrochirus*) have coexisted at this site for several decades. The bluegill population is stunted (the majority of individuals <100 mm total length), a characteristic typical of piscivore-free populations (Mittelbach 1981 and references therein). This size structure allowed minnow traps to capture most bluegill size classes (<75 mm standard length).

Twenty-four minnow traps (8-mm mesh; 40 cm long, 23 cm diameter) were placed a minimum of 5 m apart around the pond. Twice per week, traps were set on the same day during the morning (0900 h) and afternoon (1400 h). During each sampling period, half of the traps were placed in shallow, near-surface habitats (approximately 10 cm below the water surface) and half were placed in deep, near-bottom habitats at or near maximum pond depth. Traps were retrieved after two hours, and captured newts and sunfish were counted and released at the location of capture. Male and female newts were identified, whereas sunfish were not sexed due to the lack of conspicuous sexual dimorphism in juveniles and non-breeding adults.

Sampling occurred from June through May, with the exception of December through February, when the pond was completely ice-covered and inaccessible. Within each of the nine sampled months, each trap was set in each depth-by-time-of-day combination two to four times. November and March had fewer than four trapping sessions due to intermittent ice cover.

Water temperature was recorded every 30 minutes using HOBO H8 data loggers (±0.7°C; Onset Computer Corporation) deployed at multiple shallow and deep locations, but not at every trap. Temperature measurements were therefore summarized at the depth-by-time-of-day level. Sampling months were grouped into warm (June–September) and cool (October–November and March–May) seasons based on observed thermal conditions.

### Statistical analyses

#### Capture data and generalized linear mixed models

Newt and sunfish capture counts were analyzed using generalized linear mixed models (GLMMs). Counts were modeled at the trap-set level with fixed effects for season (cool vs warm), depth (deep vs shallow), time of day (morning vs afternoon), and all interactions among these factors. Trap location and sampling date were included as random intercepts to account for repeated sampling of traps and temporal non-independence among observations.

For newts, we initially explored zero-inflated negative binomial (ZINB) models to account for excess zeros. However, ZINB models that included the full interaction structure and temperature covariates exhibited non-identifiability (non–positive definite Hessians and undefined standard errors), preventing reliable inference. We therefore based inference on negative binomial (NB) GLMMs, which converged cleanly and yielded stable fixed-effect and uncertainty estimates; temperature effects were evaluated using the same NB framework to ensure comparability across models.

Sunfish counts were modeled using a NB distribution; inclusion of zero inflation was evaluated but not required for inference. Given the stability of NB estimates and consistency of conclusions across model forms, NB models were used for hypothesis testing. All mixed models were fitted using the *glmmTMB* package (Brooks *et al*. 2017) in R (version 4.4.2). Estimated marginal means (EMMs) and associated 95% confidence intervals were calculated on the response scale using the *emmeans* (Lenth 2023) package and are presented graphically to emphasize biologically interpretable contrasts.

#### Seasonal dynamics and harmonic modeling

To characterize seasonal dynamics in capture rates, we modeled newt and sunfish abundance as a function of month using harmonic regression. Harmonic models were fit to raw trap-level count data using generalized linear models with a NB error distribution and a log link. Month was treated as a circular predictor to capture continuous seasonal structure, with November and March treated as adjacent months to preserve seasonal continuity across the winter sampling gap (i.e., nine months total). Seasonal structure was modeled using sine and cosine terms of month, such that the linear predictor included sin(2πm/t) and cos(2πm/t), where *m* denotes month position on the circular scale and *t* is the total number of time units in the time series. Separate harmonic models were fit for newts and sunfish. These models were used to describe population-level seasonal trajectories and to quantify the strength and phase of seasonal cycles, but were not used for formal hypothesis testing. Depth and time-of-day effects were intentionally excluded from the harmonic models so that seasonal dynamics could be evaluated independently of finer-scale spatiotemporal structure, which was tested explicitly using the previously described mixed-effects models. Model coefficients for harmonic terms are reported to support inference about seasonal timing and amplitude.

#### Temperature data and thermal modeling

Seasonal temperature dynamics were summarized using harmonic regression models fit to monthly mean temperatures. Harmonic models included sine and cosine terms of month and were fit separately for each depth-by-time-of-day combination using ordinary least squares regression. These models were used for descriptive visualization only and were not intended for formal inference (Fig. S1).

To assess associations between temperature and capture rates independent of coarse seasonal differences, temperature was centered within season (warm and cool). Centered temperature values therefore represent deviations from the seasonal mean, isolating short-term thermal variation from broader seasonal trends. Mixed-effects models therefore included fixed effects of season, depth, time of day, centered temperature, and relevant interactions, with trap location and sampling date included as random intercepts.

Model-based relationships between temperature and capture rates were visualized using partial effect plots generated from fitted mixed-effects NB models. Predicted values and 95% confidence intervals were calculated on the response scale using the full fixed-effect variance–covariance matrix.

#### Niche overlap analysis

Niche overlap between newts and sunfish was quantified using Pianka’s index (Pianka 1973), calculated across discrete habitat–time states defined by season-by-depth-by-time of day. For each species, capture totals were summed across trap sets within each season-by-depth-by-time state and converted to proportional resource use prior to calculating overlap. Pianka’s index was calculated using a custom R function implementing the standard formulation (Pianka 1973). Null distributions were generated using 999 permutations in which sunfish capture totals were randomly reassigned among habitat–time states without replacement while preserving overall abundance. Observed overlap values were compared to null distributions to assess departures from random habitat use. Analyses were conducted for the full dataset and separately within seasons to evaluate seasonal shifts in niche overlap.

#### Sex composition of captured newts

To evaluate whether sex composition of captured eastern red-spotted newts varied across habitat and diel conditions, we analyzed sex composition conditional on capture rather than modeling sex-specific abundance. This approach directly tests whether males and females differ in habitat or diel use independent of overall activity or capture probability. Sex composition was modeled using a binomial generalized linear mixed model with a logit link, treating male and female counts per trap as a two-category response (using the *cbind* function). Fixed effects included depth, time of day, and their interaction, with trap location and sampling date included as random intercepts. Model-based estimated marginal means were calculated on the response (probability) scale using the *emmeans* package (Lenth 2023), yielding predicted probabilities that a captured newt was male for each depth-by-time combination with associated 95% confidence intervals.

Because no females were captured during the warm season, sex-specific analyses were necessarily restricted to the cool season. Under these conditions, sex-by-season interactions or sex-specific abundance models are not identifiable due to complete separation and structural zeros. Focusing on sex composition among captures during the cool season therefore represents the only statistically valid approach for assessing sex-specific habitat and diel patterns in this dataset.

## RESULTS

### Seasonal and diel patterns of capture rates

Across the annual sampling period, newts and sunfish exhibited strongly contrasting seasonal patterns in capture rates (Fig. 1). Newt captures were highest during cooler months and declined sharply during the warm season, whereas sunfish capture rates increased during warmer months and peaked in summer. Harmonic negative binomial models captured these opposing seasonal trajectories, with highly significant sinusoidal components for both taxa but opposite phase structure (Table S3), resulting in limited temporal overlap during periods of peak activity.

**Figure 1.**
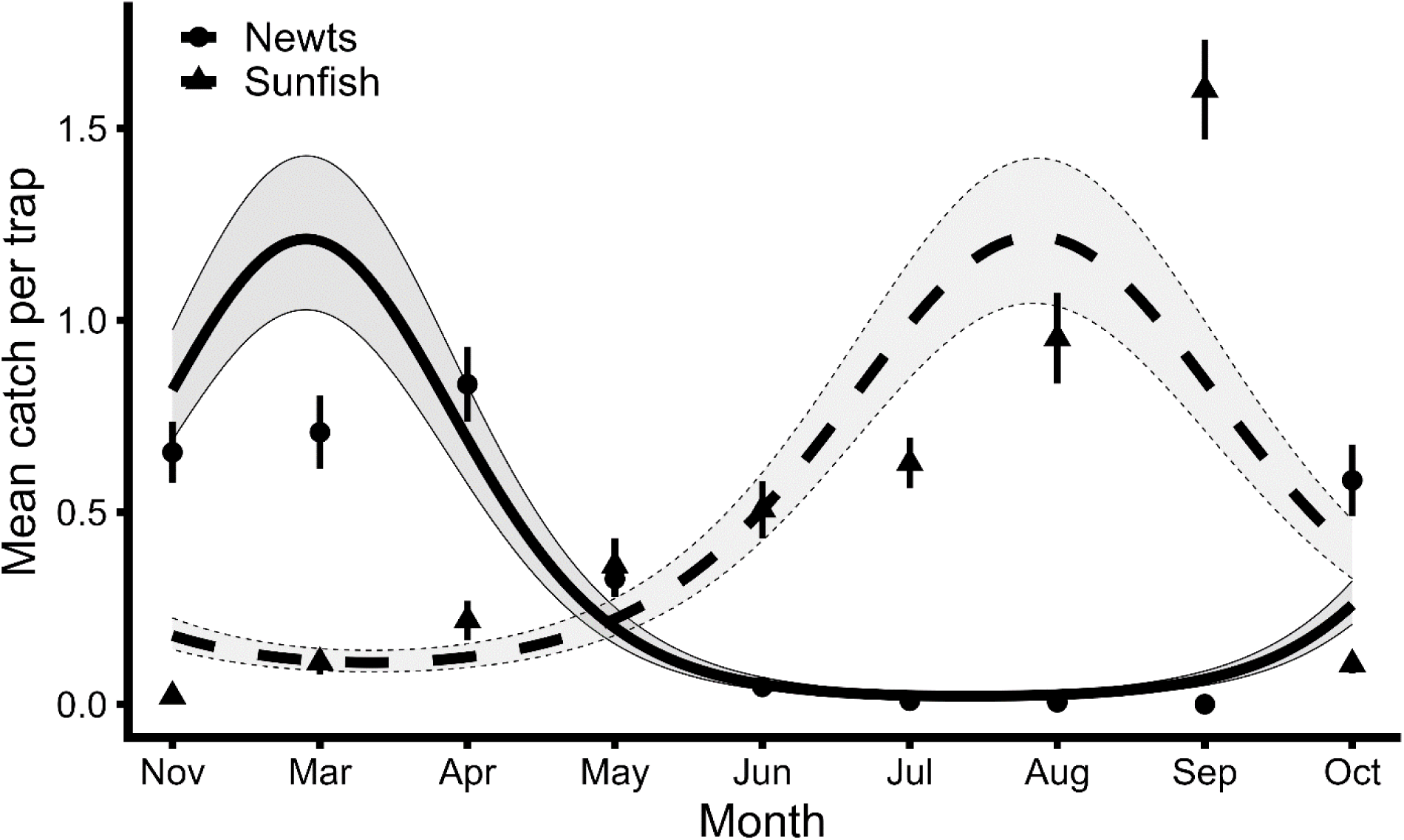
Seasonal dynamics of newt and sunfish capture rates. Points show observed monthly mean capture rates (± SE) across all traps. Solid lines show fitted values from harmonic negative binomial models, with shaded regions indicating 95% confidence intervals. Harmonic model coefficients supporting seasonal phase and amplitude are reported in Table S3.

Within seasons, the two species also differed consistently in depth and diel patterns of capture (Fig. 2). Newt captures were highest in deep habitats and during morning sampling periods, particularly during the cool season (Fig. 2A). In contrast, sunfish captures were highest in shallow habitats and tended to be higher during afternoon sampling periods, with these patterns most pronounced during the warm season (Fig. 2B). These opposing depth–diel associations indicate strong spatiotemporal partitioning that varies seasonally rather than static habitat preferences.

**Figure 2.**
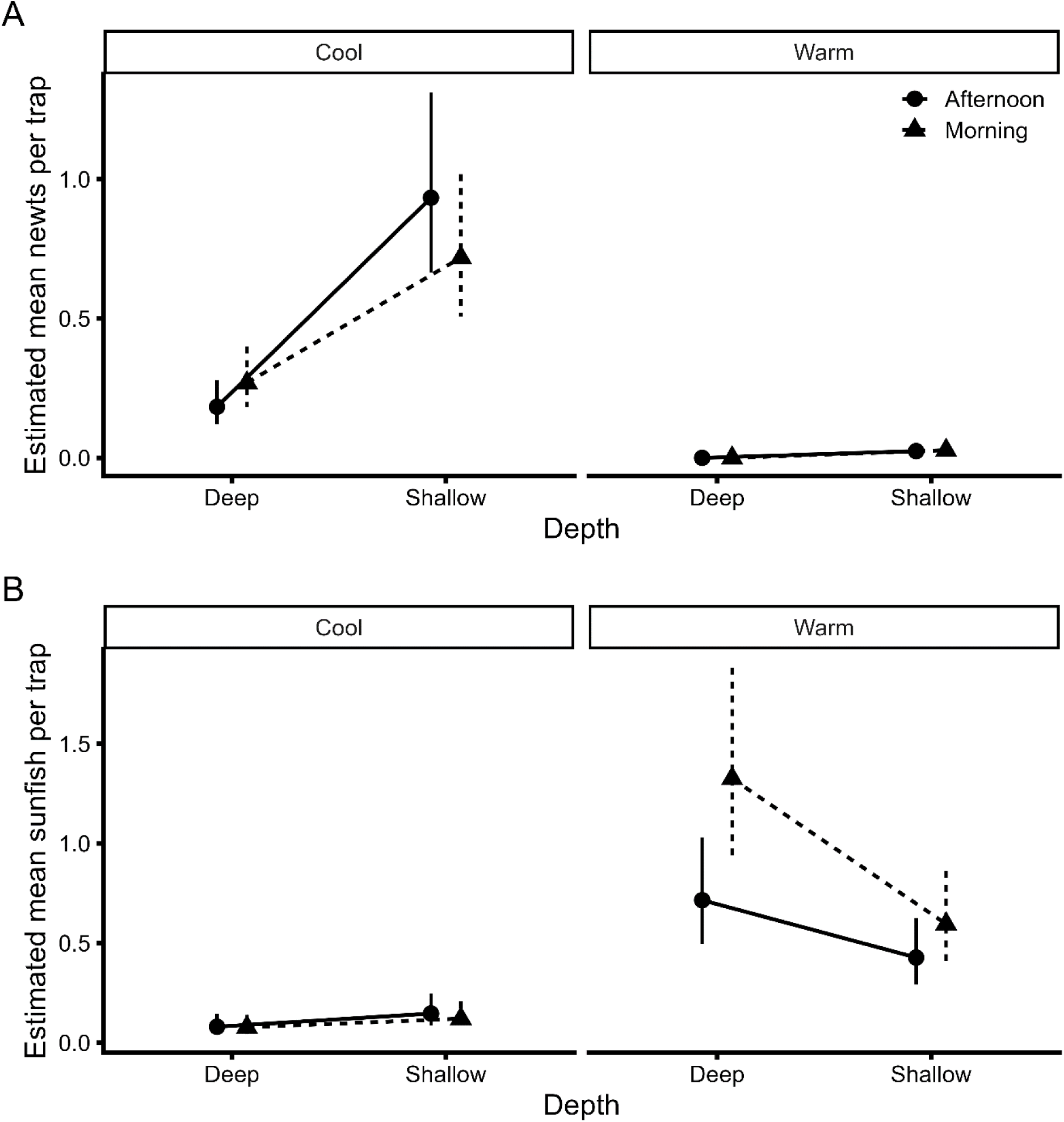
Model-based estimated marginal means (± 95% CI) of capture rates for (A) newts and (B) sunfish across depth (deep vs shallow), time of day (morning vs afternoon), and season (cool vs warm). Estimates are derived from negative binomial mixed-effects models. Formal hypothesis tests corresponding to these contrasts are reported in Table S4.

**Figure 3.**
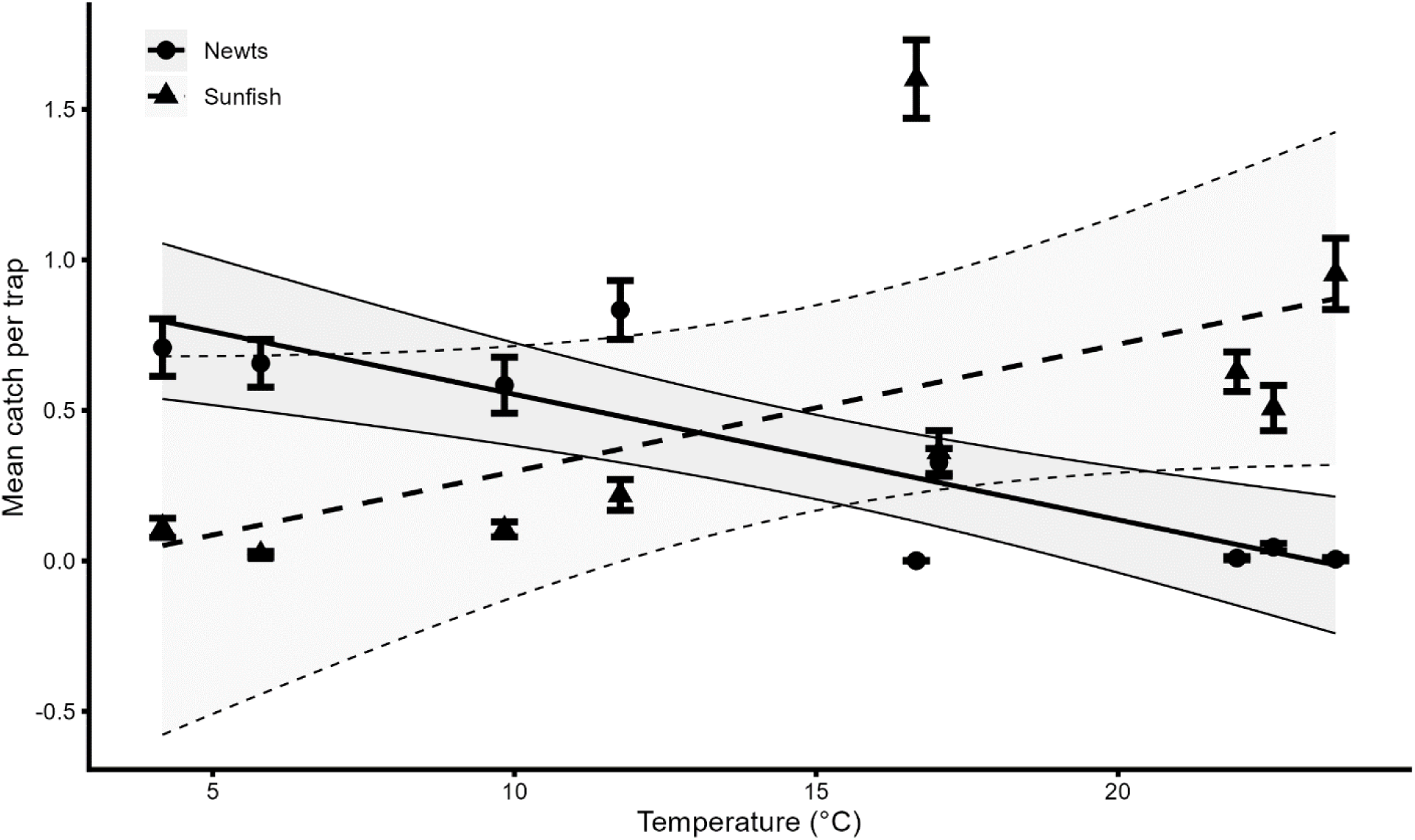
Relationships between temperature and capture rates for newts and sunfish after accounting for season, depth, and time of day. Temperature values represent deviations from seasonal means. Lines show model-based predictions from mixed-effects models, with shaded regions indicating 95% confidence intervals.

### Model-based tests of habitat, diel, and temperature effects

Negative binomial generalized linear mixed models confirmed strong and contrasting habitat associations for newts and sunfish. For newts, the negative binomial GLMM including within-season centered temperature revealed a strong effect of depth (Wald χ²□ = 79.01, *p* < 2.2 × 10□^1^□) and a significant depth-by-time-of-day interaction (Wald χ²□= 7.04, *p* = 0.00795; Table S4). On the model scale, shallow traps yielded higher captures than deep traps (β = 1.64 ± 0.18 SE), and the morning–afternoon contrast depended on depth (β = −0.66 ± 0.25 SE). Time of day alone was marginally associated with newt captures (Wald χ²□= 3.12, *p* = 0.078).

After accounting for season, depth, and time of day, deviations from the seasonal mean temperature were not associated with newt capture rates (Temp_cSeason: Wald χ² = 0.97, *p* = 0.324). Season and its higher-order interactions were not significant (all *p* ≥ 0.99), reflecting the presence of near-structural zeros in warm-season strata. Consequently, inference for newts focuses on well-estimated depth and depth-by-time effects rather than unstable season-specific coefficients.

For sunfish, NB GLMMs revealed strong seasonal increases in capture rates (Wald χ²□= 39.77, *p* < 0.001), significantly higher captures in shallow habitats (Wald χ²□= 3.96, *p* = 0.046), and a pronounced season-by-depth interaction (Wald χ²□= 10.32, *p* = 0.001; Table S5). This interaction reflected a particularly strong shift toward shallow habitat use during the warm season. Time of day did not independently influence sunfish captures (Wald χ²□= 0.01, *p* = 0.91), although afternoon-biased captures were evident in model-based predictions (Fig. 2B).

### Seasonal niche overlap between newts and sunfish

Overall niche overlap between newts and sunfish, quantified using Pianka’s index across combined season-by-depth-by-time states, was significantly lower than expected under a null model of random habitat use (O = 0.145; *p* = 0.012; Table S1; Fig. S2). When analyzed separately by season, overlap patterns diverged markedly. During the cool season, observed overlap was extremely high (O = 0.965) and exceeded null expectations, reflecting shared use of deep, morning habitats when sunfish were rare. In contrast, during the warm season, observed overlap was moderate (O = 0.465) and did not differ from null expectations (*p* = 0.181), consistent with strong spatial and diel segregation between taxa. These results indicate that temporal shifts in habitat use across the annual cycle, rather than static habitat preferences within seasons, structure niche overlap between newts and sunfish.

### Sex composition of captured newts

Sex composition of captured newts varied predictably across habitat and diel conditions during the cool season, when both sexes were present in captures. Because no females were captured during the warm season, sex-specific analyses were necessarily restricted to the cool season. A binomial mixed-effects model revealed a significant depth-by-time-of-day interaction (Wald χ² = 4.07, *p* = 0.044; Table S2). Model-based estimated marginal means indicated that morning captures in deep habitats were less male-biased than afternoon captures, whereas shallow habitats exhibited consistently high male bias with little diel variation.

## DISCUSSION

We found strong and consistent spatiotemporal niche partitioning between coexisting newts and sunfish across diel, seasonal, and habitat axes. Capture rates of the two species exhibited opposing seasonal dynamics, contrasting depth and time-of-day use, and divergent responses to temperature. These patterns resulted in low niche overlap across combined season-by-depth-by-time states relative to null expectations, suggesting that these two species may play keystone roles at different times and in different microhabitats in pond systems. Together, our results indicate that spatiotemporal segregation is a prominent feature of habitat use in this system and is consistent with stabilizing niche differences predicted by coexistence theory (Chesson 2000; Barabás, D’Andrea & Stump 2018; Ellner *et al*. 2019), even though causal links to coexistence were not directly tested.

Importantly, many of the individual components of this niche structure, such as seasonal phenologies, habitat preferences, and activity rhythms, have been documented previously when newts and sunfish were studied in isolation. Studies on newts (Hurlbert 1969; Harris, Alford & Wilbur 1988; Kesler & Munns 1991; Rohr, Madison & Sullivan 2002; Rohr, Madison & Sullivan 2003; Rohr *et al*. 2005) and on sunfish (Werner 1969; Beitinger 1975; Sarker 1977; Mittelbach 1981; Crowder & Cooper 1982; Butler 1989) independently identified patterns that closely match those observed here. Our contribution is demonstrating that these species-specific patterns align across multiple spatial and temporal axes within the same system where they cohabit, generating pronounced spatiotemporal niche partitioning and low realized overlap. This integration suggests that similar multidimensional niche division may occur in other ponds where these species coexist, a hypothesis that warrants broader comparative testing.

### Spatiotemporal structure of habitat use

Newts and sunfish displayed markedly different patterns of habitat use across seasons and diel periods. Newt capture rates were highest during cooler months, in deeper habitats, and during morning periods, whereas sunfish capture rates were highest during warmer months, in shallow habitats, and during afternoon periods. These patterns are consistent with previous work showing that bluegill and other centrarchids exhibit increased activity and foraging efficiency in warm, shallow habitats, particularly during periods of high light availability (Mittelbach 1981; Werner & Hall 1988). In contrast, adult newts may experience increased physiological constraints or altered risk–reward tradeoffs in shallow habitats during warm conditions, leading to greater use of deeper or cooler habitats and reduced surface activity (Morin 1983; Bristow 1991).

Together, these results indicate that habitat use by both species is structured by the interaction of season, depth, and time of day, producing sharply contrasting realized niches despite spatial overlap at the scale of the pond. These opposing activity patterns set the stage for pronounced spatiotemporal niche differentiation (Carscadden *et al*. 2020), which we explore further below in relation to temperature, potential mechanisms, and broader implications for coexistence.

### Temperature as a mediating factor in niche differentiation

The opposing seasonal, diel, and habitat-use patterns observed for newts and sunfish are consistent with well-established differences in their thermal biology and behavioral responses to temperature gradients. For bluegill sunfish, capture rates increased during warmer months and were highest in shallow habitats and afternoon periods, particularly during the warm season. These spatiotemporal patterns closely tracked periods and locations of elevated water temperatures and align with laboratory evidence for behavioral thermoregulation in bluegills (Magnuson & Chipman 1972; Beitinger 1975). Thus, shifts toward shallow, afternoon habitats during cooler months likely reflect active tracking of thermal maxima rather than static habitat preferences.

In contrast, eastern red-spotted newts exhibited sharply reduced capture rates during the warmest months, coincident with water temperatures approaching or exceeding reported upper thermal limits for adult *Notophthalmus* (Hutchison 1961). Although seasonal emigration of newts from ponds has often been attributed to hydrological change or pond drying (Hurlbert 1969; Harris, Alford & Wilbur 1988), our results support alternative, non-mutually exclusive explanations: that summer water temperatures become physiologically stressful or intolerable, or that increased sunfish activity intensifies competitive pressures in shallow habitats. Supporting this interpretation, closely related red-bellied newts (*Taricha rivularis*) experience physiological stress and initiate transitions toward terrestrial phases at temperatures as low as 23 °C (Licht & Brown 1967), well below peak summer temperatures observed in Harpur Pond.

At the cold end of the thermal spectrum, newts also appeared to avoid near-freezing conditions. During the coldest months, shallow morning temperatures frequently fell to approximately 3 °C or lower, and newts correspondingly shifted their morning distribution toward deeper, warmer waters. This pattern suggests fine-scale behavioral thermoregulation and aligns with experimental and field studies demonstrating active thermal habitat selection by salamanders (Skelly & Freidenburg 2000; Freidenburg & Skelly 2004; Sauer, Sperry & Rohr 2016; Cohen *et al*. 2017; Sauer *et al*. 2019).

Together, these results suggest that temperature may be acting as a powerful mechanistic mediator of spatiotemporal niche differentiation between newts and sunfish. Rather than reflecting fixed habitat preferences, the observed patterns are consistent with dynamic, behaviorally mediated responses to thermal gradients operating across seasons, depths, and diel cycles.

### Mechanisms underlying spatiotemporal niche partitioning

Several non-mutually exclusive mechanisms could underlie the pronounced spatiotemporal niche partitioning observed between newts and sunfish. Direct behavioral avoidance of the other species on ecological timescales appears unlikely because both species readily enter minnow traps baited with the other, and adult newt feeding is not reduced in the presence of bluegills (Kesler & Munns 1991; Rohr & Madison 2001; Rohr, Madison & Sullivan 2003). These findings suggest that short-term avoidance alone is an unlikely explanation for the observed patterns.

Avoidance or niche divergence on evolutionary timescales, however, remains plausible. Persistent competition can select for niche differentiation even when contemporary interactions are weak (Connell 1980; Carscadden *et al*. 2020). Experimental evidence supports this possibility; in warm-water mesocosms, newts were competitively inferior to sunfish (Bristow 1991). Under such conditions, newts may experience a competitive release by foraging during colder seasons or in microhabitats where bluegill predation pressure on invertebrates is reduced and prey populations rebound (Gilinsky 1984; Butler 1989). Selection under these conditions could favor divergence in thermal preferences, activity patterns, or both. Alternatively, differences in thermal physiology and behavioral thermoregulation may independently constrain when and where each species can forage efficiently, with niche partitioning emerging as a byproduct of physiological specialization rather than direct biotic interactions. Distinguishing between these alternatives will require experimental comparisons of activity and habitat use in the absence of the other species.

Regardless of the underlying cause, the observed partitioning is expected to reduce the strength of exploitative competition between these predators. Low spatiotemporal overlap is generally associated with weak interaction strengths (Schoener 1974), even when species share similar prey resources. The degree of segregation observed here suggests that newts and sunfish may exert strong, but temporally and spatially offset, effects on pond food webs, consistent with context-dependent keystone effects (Power *et al*. 1996).

### Niche overlap and implications for coexistence

Observed niche overlap between newts and sunfish was low across combined season-by-depth-by-time states and consistently lower than expected under null models of random habitat and time use. Low overlap across multiple niche dimensions is consistent with stabilizing niche differences that can reduce interspecific interactions relative to intraspecific interactions (Chesson 2000; Levine & HilleRisLambers 2009; Barabás, D’Andrea & Stump 2018; Ellner *et al*. 2019).

However, our results do not demonstrate that spatiotemporal niche partitioning causes coexistence in this system. Experimental manipulations or demographic data would be required to establish causal links between niche differentiation and population persistence. Instead, our findings quantify the structure and strength of niche partitioning among coexisting top predators and demonstrate that their realized niches overlap less than expected by chance—a necessary, though not sufficient, condition for coexistence mediated by niche differences.

### The importance of scale in detecting spatiotemporal niche partitioning

This study highlights the critical importance of scale in detecting and interpreting niche partitioning. The synchronous but opposing shifts in habitat use and diel activity by newts and sunfish would have been underestimated, or missed entirely, had sampling been conducted at only a single temporal or spatial scale. Partitioning was evident only when capture data were analyzed across multiple nested scales, including within days, across microhabitats, and over seasonal cycles.

Paradoxically, niche separation may have been least apparent during periods when newts and sunfish briefly overlapped while transitioning between habitats or diel activity regimes. A study conducted solely during these transition periods could have erroneously concluded that niche overlap was high and partitioning weak. More broadly, many studies of diel activity are conducted at a single time of year, and many seasonal studies implicitly average over diel variation, potentially masking scale-dependent patterns (Power 2001; Rohr, Madison & Sullivan 2003; Cohen *et al*. 2016; Nakabayashi *et al*. 2021; Bloomfield *et al*. 2022).

### Broader implications and limitations

The newt–sunfish system represents a relatively rare case of an amphibian species coexisting with fish, a context in which amphibians often experience strong negative effects (Kats, Petranka & Sih 1988; Wellborn, Skelly & Werner 1996). Although eastern red-spotted newts possess tetrodotoxin across life stages (Brodie 1968; Brodie Jr, Hensel Jr & Johnson 1974), toxicity alone does not preclude interactions with fish. Our results indicate that spatiotemporal niche partitioning may complement chemical defenses by reducing overlap during periods or in habitats where interactions are most intense.

Several limitations warrant consideration. Capture rates provide an indirect measure of activity and habitat use and may be influenced by trap-related behaviors. Temperature measurements were summarized at the depth-by-time scale and therefore do not capture fine-scale thermal heterogeneity among individual trap locations. Despite these limitations, the consistency of patterns across analytical approaches and the concordance among seasonal, diel, and thermal results support the robustness of our conclusions.

### Conclusions

By integrating year-round trapping data, hierarchical count models, temperature-informed analyses, and null-model tests of niche overlap, this study provides a detailed characterization of spatiotemporal niche partitioning between coexisting newts and sunfish. Our results demonstrate that niche differentiation across diel, seasonal, and habitat axes is pronounced and associated with opposing responses to temperature, underscoring the importance of multiscale approaches for understanding species interactions and coexistence in natural communities.

## ACKNOWLEDGEMENTS

This research was supported by grants from the National Science Foundation (DEB 2109293, DEB 2017785, IOS 1754868) to J.R.R.

## Supplementary Information

**Figure S1.**
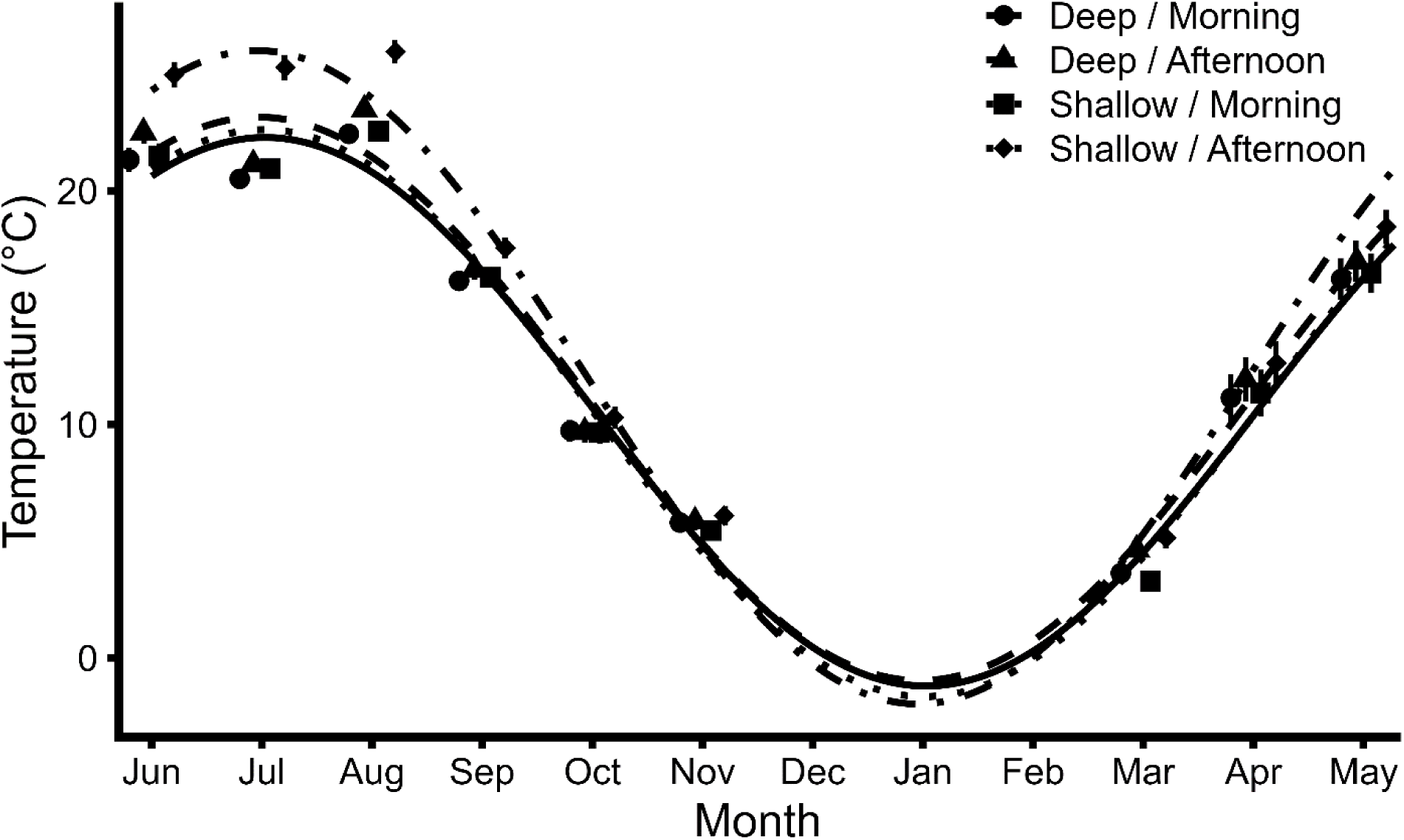
Seasonal temperature dynamics across depth-by-time-of-day strata, summarized using harmonic regression. Lines represent fitted harmonic functions; points show monthly means ± SE. These models were used for visualization only.

**Figure S2.**
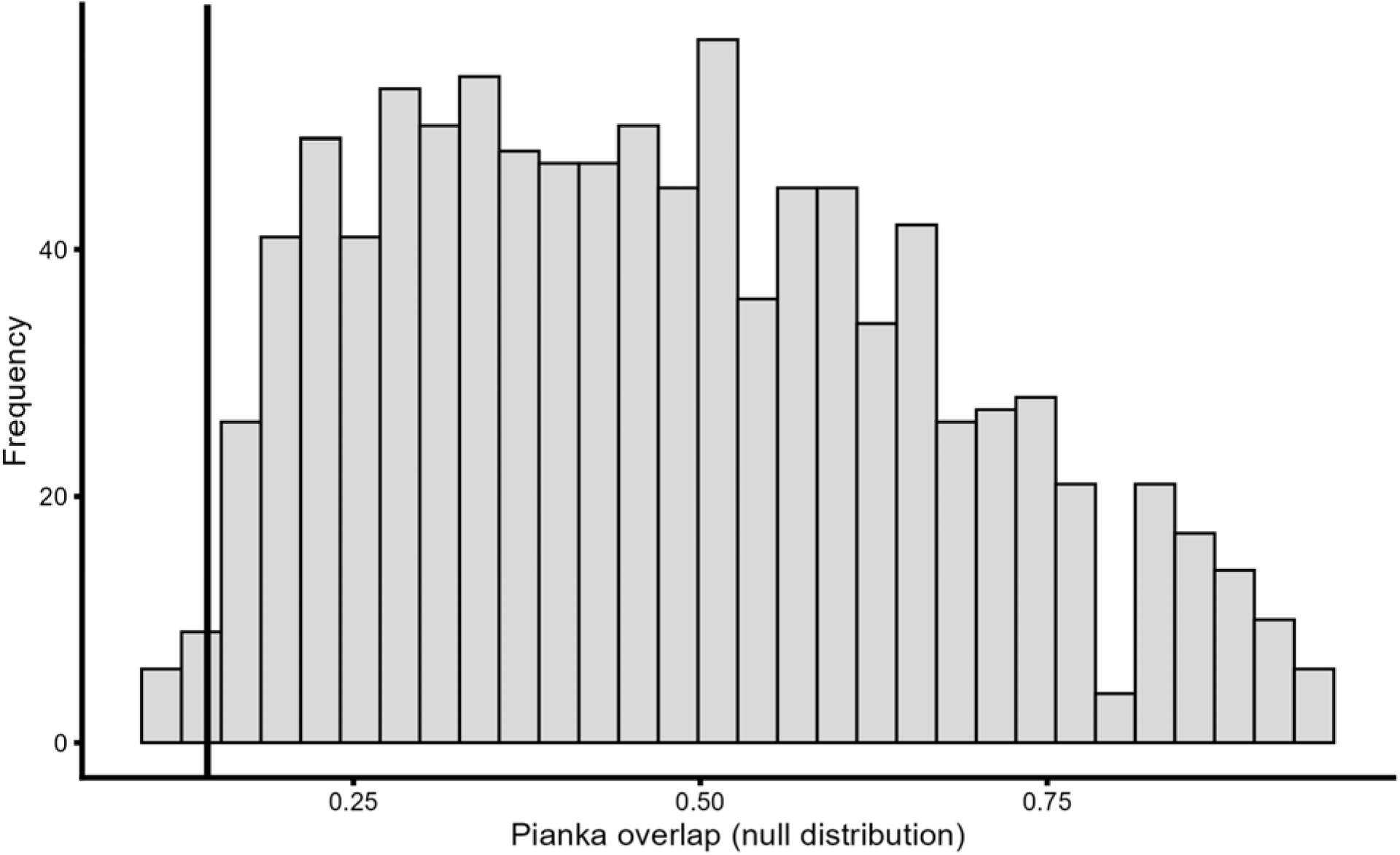
Null distribution of Pianka’s niche overlap index generated by permutation of habitat–time states. The vertical line indicates the observed overlap value. Summary statistics and permutation-based *p*-values are reported in Table S1.

**Table S1.** Niche overlap between eastern red-spotted newts (Notophthalmus viridescens) and bluegill sunfish (Lepomis macrochirus) quantified using Pianka’s index across combined season-by-depth-by-time-of-day resource states. Observed overlap (Oobs) is compared to null expectations (Onull) generated from 999 permutations that randomized sunfish capture totals among resource states while preserving marginal totals. Reported values include the mean and standard deviation (sd) of null overlap distributions and lower-tail permutation p-values (p lower).

| Season | Oobs | Onull mean | Onull sd | p lower |
| --- | --- | --- | --- | --- |
| Overall | 0.145 | 0.473 | 0.197 | 0.012 |
| Cool season only | 0.965 | 0.837 | 0.074 | 1.000 |
| Warm season only | 0.465 | 0.650 | 0.146 | 0.181 |

**Table S2.** Model-based estimated marginal means (±95% confidence intervals) for the probability that a captured eastern red-spotted newt was male, across depth and time-of-day combinations during the cool season. Estimates are derived from a binomial generalized linear mixed model with trap location and sampling date as random intercepts.

| Depth | Time | Probability male | SE | asympt.LCL | asympt.UCL |
| --- | --- | --- | --- | --- | --- |
| Deep | Afternoon | 0.960 | 0.028 | 0.848 | 0.990 |
| Shallow | Afternoon | 0.914 | 0.021 | 0.863 | 0.947 |
| Deep | Morning | 0.829 | 0.050 | 0.707 | 0.906 |
| Shallow | Morning | 0.926 | 0.021 | 0.872 | 0.958 |

**Table S3.** Coefficients from negative binomial harmonic regression models describing seasonal dynamics in newt and sunfish capture rates. Models include sine and cosine terms to capture cyclical seasonal structure across nine sampled months. Estimates are shown on the log scale.

| Term | Estimate | SE | z | p-value |
| --- | --- | --- | --- | --- |
| Newts |  |  |  |  |
| Intercept | -1.834 | 0.082 | -22.349 | 1.24E-110 |
| sin | 1.968 | 0.109 | 18.046 | 8.41E-73 |
| cos | 0.481 | 0.094 | 5.107 | 3.27E-07 |
| Sunfish |  |  |  |  |
| Intercept | -1.011 | 0.062 | -16.307 | 8.79E-60 |
| sin | -1.206 | 0.088 | -13.761 | 4.37E-43 |
| cos | 0.086 | 0.077 | 1.117 | 2.64E-01 |

**Table S4.** Type III Wald χ² tests from a negative binomial generalized linear mixed model evaluating the effects of season (cool vs. warm), depth (deep vs. shallow), time of day (morning vs. afternoon), and their interactions on eastern red-spotted newt (Notophthalmus viridescens) capture rates. Models included random intercepts for trap location and sampling date. Zero-inflated models failed to converge reliably; therefore, inference is based on the negative binomial model (see Methods).

| Effect | $\chi^2$ | df | p-value |
| --- | --- | --- | --- |
| Season | 0.000 | 1 | 0.9940 |
| Depth | 77.967 | 1 | < 0.0001 |
| Time of day | 3.455 | 1 | 0.0630 |
| Season × Depth | 0.000 | 1 | 0.9950 |
| Season × Time | 0.000 | 1 | 0.9990 |
| Depth × Time | 6.712 | 1 | 0.0096 |
| Season × Depth × Time | 0.000 | 1 | 0.9990 |

**Table S5.** Type III Wald χ² tests from a negative binomial generalized linear mixed model evaluating the effects of season (cool vs. warm), depth (deep vs. shallow), time of day (morning vs. afternoon), and their interactions on sunfish capture rates. Models included random intercepts for trap location and sampling date. Zero-inflated models provided only marginal improvement in model fit and were not required for inference; therefore, results are based on the negative binomial model.

| Effect | $\chi^2$ | df | p-value |
| --- | --- | --- | --- |
| Season | 39.774 | 1 | < 0.0001 |
| Depth | 3.965 | 1 | 0.0460 |
| Time of day | 0.012 | 1 | 0.9110 |
| Season × Depth | 10.324 | 1 | 0.0010 |
| Season × Time | 3.186 | 1 | 0.0740 |
| Depth × Time | 0.131 | 1 | 0.7180 |
| Season × Depth × Time | 0.071 | 1 | 0.7900 |

**Table S6.** Incidence rate ratios (IRRs) and 95% confidence intervals for biologically relevant contrasts in capture rates of eastern red-spotted newts (Notophthalmus viridescens) and sunfish (Lepomis spp.), derived from estimated marginal means of negative binomial generalized linear mixed models. IRRs represent multiplicative changes in expected capture rates between habitat or diel conditions and are presented to facilitate biological interpretation of model effects.

| Contrast | IRR | 95% CI | p-value |
| --- | --- | --- | --- |
| Newts |  |  |  |
| Deep vs. Shallow (Cool, Morning) | 5.12 | 3.90–6.73 | <0.001 |
| Morning vs. Afternoon (Deep) | 1.47 | 0.98–2.20 | 0.063 |
| Deep vs. Shallow (Cool, Afternoon) | 3.84 | 2.81–5.25 | <0.001 |
| Sunfish |  |  |  |
| Warm vs. Cool (Shallow) | 8.9 | 5.60–14.1 | <0.001 |
| Shallow vs. Deep (Warm) | 2.18 | 1.35–3.52 | 0.001 |
| Afternoon vs. Morning (Warm) | 1.31 | 0.98–1.76 | 0.074 |

